# Exploring the Efficiency of Deep Graph Neural Networks for RNA Secondary Structure Prediction

**DOI:** 10.1101/2024.10.11.617338

**Authors:** Md. Sharear Saon, Kevin Boehm, Grace Fu, Ian Hou, Jerry Yu, Brent M. Znosko, Jie Hou

## Abstract

Ribonucleic acid (RNA) plays a vital role in various biological processes and forms intricate secondary and tertiary structures associated with its functions. Predicting RNA secondary structures is essential for understanding the functional and regulatory roles of RNA molecules in biological processes. Traditional free-energy-based methods for predicting these structures often fail to capture complex interactions and long-range dependencies within RNA sequences. Recent advancements in machine learning, particularly with graph neural networks (GNNs), have shown promise in enhancing the ability to model the relationships between molecular sequences and their structures. This work specifically explores the efficacy of various GNN architectures in modeling RNA secondary structure. Through benchmarking the GNN methods against traditional energy-based models on standard datasets, our analysis demonstrates that GNN models improves traditional methods, offering a robust framework for accurate RNA structure prediction.

## 1. Introduction

Ribonucleic acid (RNA) is a fundamental macromolecule essential for life, on par with DNA and proteins, and plays a critical role in various biological processes, including catalysis, translation, antiviral immunity, gene expression regulation and protein synthesis [1, 2, 3, 4]. RNAs are classified into two categories: messenger RNA (mRNA), which is translated into proteins, and non-coding RNA (ncRNA), such as tRNA and rRNA, which do not encode proteins [5]. According to the central dogma of molecular biology, genetic information is transferred from DNA to RNA, subsequently guiding protein synthesis [6].

RNA molecules form complex secondary and tertiary structures from their primary sequences, which are closely tied to their biological functions. It remains challenging to determine both secondary and tertiary structures from sequence due to the limited availability of high-resolution structural data [7, 8]. Secondary structure, which influences gene expression, can be classified based on topological number, or genus, into planar structures and pseudoknots [9]. Meanwhile, tertiary structures provide precise architecture that defines enzymatic active sites and selective ligand binding pockets [10] and are primarily determined by the local conformation at secondary-structure junctions and sequence-specific interactions [10, 11].

Experimental methods, such as X-ray crystallography (X-ray), single-particle cryo-electron microscopy (cryo-EM), nuclear magnetic resonance spectroscopy (NMR) [12], can be employed to determine RNA structures. Additionally, the combination of thermodynamic and comparative sequence analyses is a widely-used technique for predicting RNA secondary structure from a single sequence. Diverse computational methods have also been developed to predict both secondary and tertiary structures from RNA sequences [13, 14]. Specifically, the prediction of secondary structure from the sequence is a crucial intermediate step in predicting tertiary structure. For instance, fragment-based methods to predict RNA tertiary structures involve breaking down the RNA’s secondary structure into its constituent elements, such as stems, hairpin loops, bulge loops, and internal loops [15, 16, 17, 18]. Subsequently, the tertiary structures of these motifs are identified by searching against a database of previously solved RNA structures. These three-dimensional fragments are then assembled to predict the full tertiary structure. Therefore, the accurate prediction of secondary structural motifs from the RNA sequence is crucial to improve the RNA tertiary structure prediction.

Several databases dedicated to RNA sequence and structure are well-maintained to facilitate RNA computational analyses. Notably, the RNAcentral database houses 35 million known RNA sequences [19]. In contrast, the Nucleic Acid Database, as of July 2024, contains only 17,934 experimentally-determined structures [20]. Research indicates that merely about 3,335 non-redundant, representative RNA 3D structures exist for structure modeling [21], with most being fragments < 100 base pairs.

In addition to these resources for primary sequences and tertiary structures, the bpRNA meta dataset [22] collects RNA secondary structures from various sources, including the Comparative RNA Web (CRW) [23], transfer messenger RNA (tmRNA) database [24], Signal Recognition Particle (SRP) Database [25], tRNAdb 2009 Database [26], RNase P (RNP) Database [27], Research Collaboratory for Structural Bioinformatics Protein Data Bank (RCSB PDB) [28], and the RNA Family Database (RFAM) [29]. Moreover, the RNA Characterization of Secondary Structure Motifs (RNA CoSSMos) database offers detailed structural features of RNA motifs extracted from previously solved RNA structures [30].

The availability of a comprehensive RNA knowledge repositories facilitate the development of various statistical and machine learning algorithms for predicting RNA secondary structures. For many decades, free energy minimization has been the predominant method for predicting RNA secondary structures. These methods leverage thermodynamic parameters to estimate the free energy changes associated with the formation of various potential secondary structures [13]. Notable energy-based methods include RNAfold [31], Centroidfold [32], Contrafold [33] and LinearPartition [34], along with machine learning based thermodynamic approaches, MXfold [35]. Moreover, the integration of deep learning techniques has significantly enhanced the analysis of RNA sequence and structural properties, offering substantial improvements over traditional comparative sequence analysis methods [36, 37, 38, 39, 40, 41].

Graph neural networks (GNNs), renowned for their efficacy in molecular structure analysis, are particularly adept at capturing the intricate connectivity patterns of RNA molecules, making them highly suitable for modeling graph-structured data and long-range interactions.

In this study, we conducted a comprehensive evaluation of several widely-used GNN architectures for RNA secondary structure prediction. We consider the RNA structure as a network, as illustrated in Figure 1(C), where each node (vertex) represents a nucleotide, and each link (edge) denotes a base pair between nucleotides. We selected six GNN methods that demonstrated robust performance: graph convolutional networks (GCNConv) [42], graph attention networks (GATConv) [43], gated graph convolution(GatedGNN) [44], edge-Conditioned Convolutions (EdgeConv) [45], auto-regressive moving average graph convolution (ARMAConv) [46] and graph neural networks with personalized pagerank (APPNPConv) [47]. Our methods were trained and evaluated using a well-established RNA dataset, including bpRNA and PDB datasets for pre-training and performance validation. The code and data associated with this study are publicly accessible at https://github.com/JieHou-SLU/RNASS.

**Figure 1.**
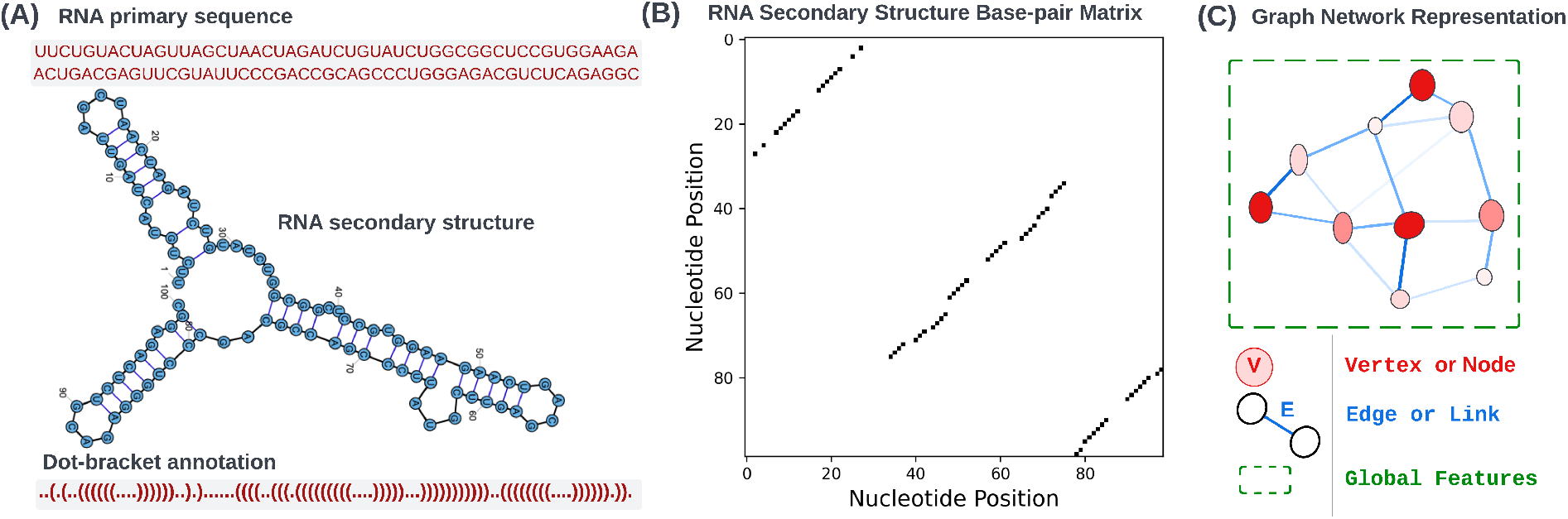
Multimodal Representations of RNA Secondary Structure. (A) RNA sequence with its corresponding dot-bracket annotation and visual secondary structure; (B) a binary base-pair matrix illustrating nucleotide interactions; (C) presents a graph network visualization of the RNA structure, highlighting the connections between nucleotides and how information is aggregated from neighboring nodes to form global features using GNN.

## 2. Related work

Our work extends current state-of-the-art deep learning models for RNA secondary structure prediction by exploring the potential application of graph neural networks. Several state-of-the-art deep learning methods process RNA nucleotide sequences using either one-dimensional feature data or two-dimensional, image-like formats suitable for convolutional neural networks (CNNs) and similar architectures [36, 37, 38, 39, 40, 41]. These approaches facilitate the learning of long-range interactions in RNA secondary structures, which are typically represented as simple strings in dot-bracket notation or as nucleotide base-pair maps. For instance, Figure 1(A) displays the RNA sequence along with its corresponding secondary structure in dot-bracket annotation, where dots represent unpaired nucleotides and parentheses denote base pairs. Figure 1(B) shows the binary basepair contact map between nucleotides in sequence. Each dot in map represents a base pair between two nucleotides, plotted according to their positions in the sequence.

The method DMfold represents the first integration of deep learning with an enhanced base pair maximization principle for RNA secondary structure prediction that includes pseudoknots [38]. This method automates the calculation of folding parameters through deep learning, requiring only the target sequence as input, thereby simplifying operational procedures.

Another method, named SPOT-RNA, utilizes deep neural network learning techniques for RNA secondary structure prediction, encompassing both local and nonlocal interactions [37]. It demonstrates superior performance compared to existing algorithms, particularly in predicting noncanonical base pairs, though it shows comparatively lower efficacy in pseudoknot predictions, achieving the highest F1-score of 0.239. Notably, its precision rate achieves 13% improvement over the subsequent leading method, MX-fold [35]. SPOT-RNA2, an advancement over its predecessor, enhances prediction accuracy by incorporating sequence profiles and direct mutational coupling. This enhancement extends to canonical base pairs and complex tertiary interactions, including non-canonical pairs and pseudoknots [39]. Its ensemble deep neural network architecture, inspired by both SPOT-RNA and RNAsnap2 [48], attains an F1-score > 0.8 for RNAs with > 1000 homologous sequences.

CDPfold introduces a novel approach by merging convolutional neural networks with dynamic programming for RNA secondary structure prediction [36]. It leverages CNNs to extract latent features from RNA sequence data and applies dynamic programming to refine these predictions, showing notable improvements, especially in datasets lacking pseudoknots, with a 30% higher success rate than comparable algorithms.

E2Efold, an end-to-end deep learning model, incorporates hard constraints into its architecture for RNA secondary structure prediction [49]. It excels in predicting pseudoknotted structures and maintains efficiency, outperforming many state-of-the-art algorithms in inference time.

MXfold2, building upon the foundation set by MXfold [35], integrates thermodynamic regularization with Turner’s nearest neighbor free energy parameters in its deep neural network training process to enhance RNA secondary structure prediction accuracy [40]. Experimental evidence suggests that this thermodynamic integration substantially boosts prediction accuracy, with MXfold2 achieving superior results on unseen dataset without suffering overfitting on training set.

Lastly, UFold adopts a new deep learning architecture, named U-Net, to predict RNA secondary structures. It transforms nucleotide sequences into an image-like format for training on annotated RNA base-pair datasets. UFold’s predictions are rapid, with an inference time of 160 ms per sequence for lengths up to 1500 base pairs [41].

Traditional sequence-based deep learning methods with CNN architecture often struggle with long-range dependencies due to their sequential processing nature. In contrast, GNNs excel at capturing long-range interactions through a series of graph aggregation layers. These layers propagate node information across both local and global neighbors within the graph structure, allowing interactions between distantly located nucleotides to directly influence the prediction. Recently, GNNs have emerged as a powerful tool for learning representations of bioinformatics data, particularly excelling in molecular structure predictions and chemical property predictions where data inherently form graph structures [50, 51, 52, 53, 54]. For example, a protein’s 3D structure can be transformed into a graph representation, enabling the learning of dependencies among amino acids in three-dimensional space. In this graph representation, all non-hydrogen atoms of the protein serve as nodes, while the chemical bonds connecting these atoms in the 3D model act as edges. Thus, the graph representation encodes both the sequence and structural information, facilitating the exploration of relationships between sequence information (node-level embedding) and spatial proximity (edge-level embedding). In the context of RNA secondary structure prediction, GNNs model the intricate relationships between nucleotide sequences and their secondary structures. By combining local and global contextual information, GNNs substantially improve the accuracy of structure predictions.

In this paper, we introduce a GNN framework tailored for RNA secondary structure prediction. We explore the efficacy of various GNN training strategies applied to RNA datasets and demonstrate how integrating energy-based methods with GNNs can significantly enhance performance in predicting RNA secondary structures. Furthermore, we highlight potential future directions and improvements to enhance the performance of GNNs in the prediction of RNA secondary or tertiary structures.

## 3. Materials and methods

### 3.1 Data Preparation

To assess the capability of graph neural networks for RNA secondary structure prediction, we utilized datasets curated in prior studies [37, 39] for model training, validation, and testing. Specifically, we employed the first version of the bpRNA-1m dataset, which comprises 10,714 sequences for training (TR0), 1,274 for validation (VL0), and 1,266 for testing (TS0). Each of these datasets features RNA sequences alongside their annotated secondary structures. Given that many secondary structures in the bpRNA-1m database are computationally predicted rather than experimentally validated, we also evaluated the thermodynamic stability of these structures using ‘efn2’, a tool from the RNAstructure package [13]. Analysis of the free energy across the bpRNA datasets, as depicted in Figure 2, reveals that 11.9% of the structures exhibit non-negative free energy values, indicating thermodynamically unfavorable configurations. These include less stable local structures such as small loops with fewer than three nucleotides and short stems with weaker base pairs (e.g., GU wobbles). To enhance the reliability of our dataset, we refined the bpRNA dataset by excluding RNA sequences whose annotated secondary structures exhibited unfavorable free energy values, setting a cutoff value of -10 kcal/mol based on preliminary analyses. Additionally, due to hardware constraints, the maximum sequence length for model training was capped at 500. This filtering and limitation were adopted to optimize the training process and improve the predictive accuracy of the GNN models.

**Figure 2.**
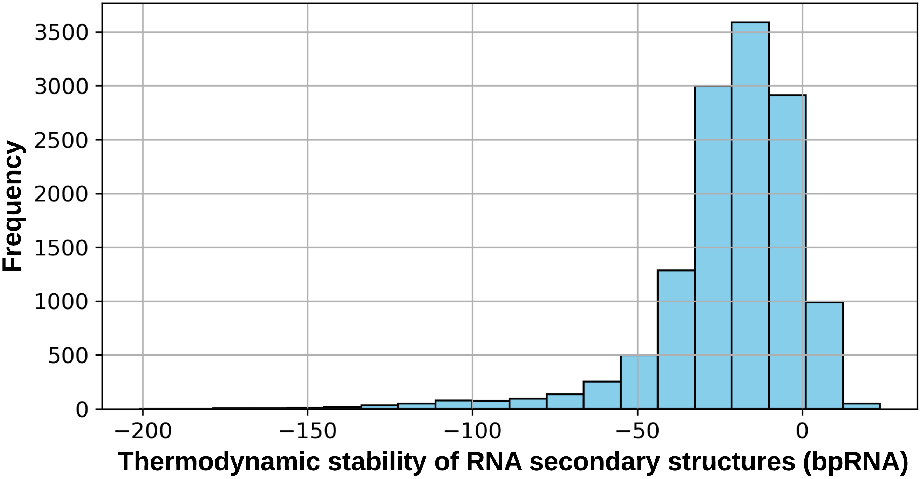
Distribution of thermodynamic stability of RNA secondary structures in bpRNA dataset calculated based on ‘efn2’ in RNAstructure package [13]

In addition to the bpRNA datasets, we also applied transfer learning to fine-tune our GNN models using RNA secondary structures derived from experimentally verified structures in the PDB database. As detailed in prior research [37, 39], the PDB dataset includes 120 sequences for training (TR1), 30 for validation (VL1), 66 for independent testing (TS1) and a rigorous blind test set, designated as TS-hard, comprising 23 RNA sequences that exhibit minimal or no similarity to those in the training and validation sets. This hard set is specifically designed to evaluate the model’s ability to generalize across unfamiliar RNA families.

### 3.2. Feature Generation

This section outlines the node and edge features used for graph learning, as depicted in Figure 3.

**Figure 3.**
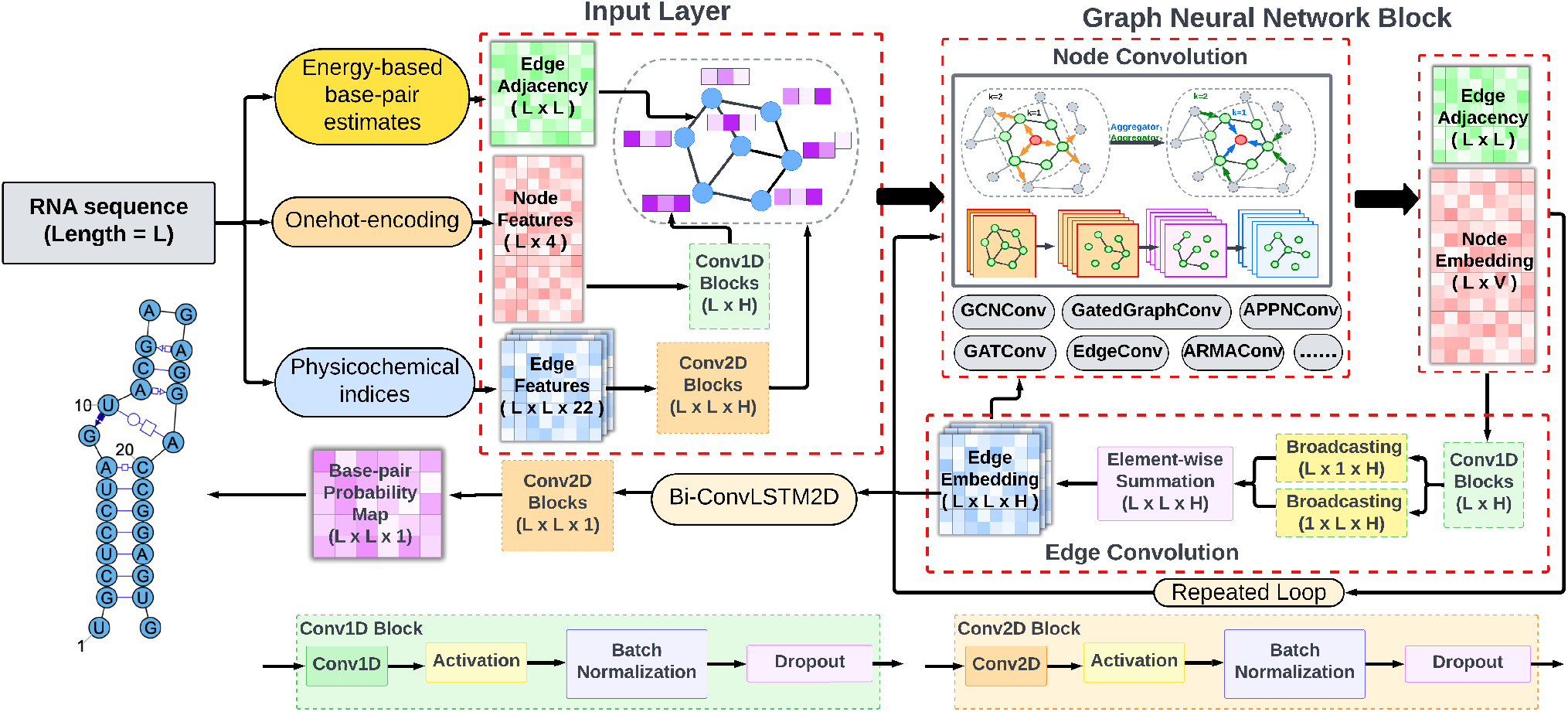
Proposed graph neural network framework for RNA secondary structure prediction: This framework integrates energy-based base-pair predictions with diverse graph learning methods. (A) Input layer: utilizes RNA sequence data alongside physicochemical indices and energy-based base-pair estimates to construct node and edge features. (B) Graph neural network block: employs repeated execution of various graph convolution layers (GCNConv, GatedGraphConv, GATConv, EdgeConv, APPNPConv, ARMAConv) to refine node and edge features. (C) Edge embedding and convolution: performs edge convolution to enhance RNA interaction representations, followed by a bi-directional ConvLSTM2D layer that captures spatial dependencies and generate base-pair probability predictions.

#### 3.2.1. Node-level representations for GNNs

For an RNA sequence input, each nucleotide is treated as an independent node within an undirected graph. The feature representation for each node employs one-hot encoding, where the four nucleotide bases (A, U, C, G) are each converted into a binary vector of length four, distinctly representing the nucleotides. In this format, each position in the vector corresponds to a nucleotide; the position representing the nucleotide present in the sequence is marked with a 1, while all other positions are marked with a 0. This approach generates a node feature matrix of shape (L x 4) for a sequence with length L. The node features are transformed via a 1D convolution block, which extracts hidden features, commonly referred to as embeddings. This convolution block consists of a series of layers: a 1D convolution layer equipped with six filters and a kernel size of five, an activation layer employing rectified linear units (ReLU) [55], followed by a batch normalization layer [56]to generate the node embeddings.

#### 3.2.2. Edge-level representations for GNNs

To predict pairwise nucleotide interactions in RNA, we utilize two matrices as inputs for GNNs: the nucleotide-nucleotide adjacency matrix and the nucleotide-nucleotide feature matrix, both with dimensions (L x L x K). The adjacency matrix, a binary matrix denoted as *A*, encapsulates potential interactions between nucleotides, where a value of 1 indicates a direct interaction (e.g., hydrogen bond or base pairing) between nucleotides *i* and *j*, and a 0 denotes no interaction. Initial connectivity between nucleotides is determined using base-pair probabilities predicted by energy-based predictors, including LinearPartition [34], and thermodynamic method, MXfold [35]. Both methods utilize thermodynamic models to assess the likelihood of each nucleotide pair being bonded in the native structure. Pairs with a probability > 0.5 are considered likely to form a stem in the RNA secondary structure. It is worth noting that other free-energy-based methods for estimating the stability of RNA structures and base-pairs can be adopted as initial connectivity for graph learning.

GNNs facilitate feature learning across all graph edges, defined by the adjacency matrix. In our study, we included 22 physicochemical indices for all 16 dinucleotides, detailing the physical and chemical properties of RNA nucleotides and their interactions, as specified in Pse-in-One 2.0 [57]. These indices include both content information—such as adenine and cytosine content, GC content, guanine content, and others related to structural attributes like enthalpy, entropy, free energy, and stacking energy—and dynamic properties like hydrophilicity and molecular movements (rise, roll, shift, slide, tilt, and twist). Consequently, the feature matrix representing these RNA physicochemical properties is structured as (L x L x 22), providing a comprehensive set of edge features for GNN processing. This approach allows for a nuanced understanding of RNA interactions, crucial for accurate secondary structure prediction.

We use the Embedding layer provided in Keras [58] to convert the binary integers in adjacency matrix into dense vectors with output dimension of 20 as a part of edge representation. We also add one 2D convolution block layer with 20 channels on top of the physicochemical features to map the physicochemical indices into dense vectors as the second part of edge representation.

### 3.3. Graph Neural Network Training

We adopted a sophisticated graph-based neural network architecture similar to those described in [59, 52], but is redesigned to accommodate diverse graph convolution layers for predicting pairwise nucleotide interactions in RNA sequences. The detailed design of our architecture is depicted in Figure 3. Our model integrates multiple input representations, including node and edge features, which are processed through an advanced combination of graph convolution and recurrent layers.

#### Model Inputs and Feature Embedding

The node features begin as one-hot encoded vectors representing the four nucleotide types (A, U, C, G) in the RNA sequence.

These vectors undergo initial processing through instance normalization and a 1D convolutional layer. Edge features are represented in two forms: a binary adjacency matrix that indicates direct nucleotide interactions, and a feature matrix that contains up to 22 channels detailing physicochemical properties.

#### Graph Convolution Processing

The core of our GNN model consists of several graph convolution layers that process the node features. We evaluated and compared different types of graph convolution layers, such as GCNConv [42], GATConv [43], GatedGNN [44], EdgeConv [45], ARMAConv [46], and APPNPConv [47], based on the experiment configuration. These layers are specifically chosen to enhance the model’s capability to aggregate information from neighboring nodes, effectively capturing the complex connectivity patterns inherent in RNA structures. Additionally, other graph convolution layers, as implemented in [60], can also be integrated into this module. We experimented with various GNN architectures provided by the ‘Spektral’ package [60], and ultimately selected the six best-performing GNN layers for evaluation in this study.

As shown in Table 1, graph representations are characterized primarily by two components: node features *X*_*i*_ and edge features *E*_*ij*_. These features represent the attributes of graph nodes and the connections between the *i*^*th*^ and *j*^*th*^ nodes, respectively. The transformations applied to these node and edge embeddings through each graph convolutional block are detailed in Table 1. Here, *C*_1*D*_(.) denotes a 1D convolution block that includes a 1D convolution layer followed by batch normalization and a ReLU activation function. Similarly, *C*_2*D*_(.) encompasses a 2D convolution block comprising a 2D convolution layer, batch normalization, and a ReLU activation function. The functions *F* ^*Node*^(.) and *F* ^*Edge*^(.) are used for feature transformation through any of different graph convolution layers (e.g., GCNConv, GATConv, GatedGNN, EdgeConv, ARMAConv, APPNPConv). *X*^*l*^ and *E*^*l*^ indicate the transformations of node and edge features, respectively, at the *l*^*th*^ layer in the graph learning process.

**Table 1.**
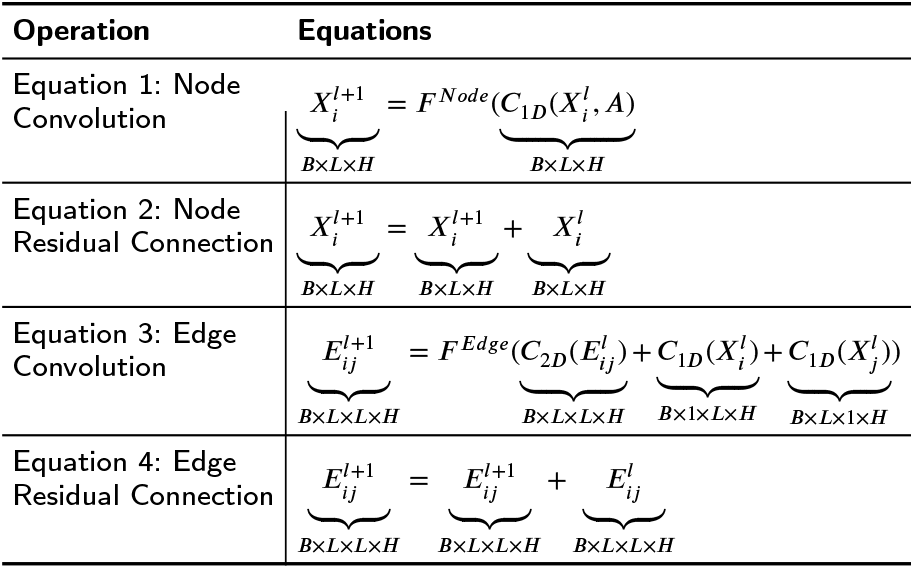
Node and edge operations in graph convolution neural network.

In the node convolution block (referenced as Equation 1 in Table 1), node embeddings are processed using 1D convolutional filters. The convolved outputs are then fed into a designated node convolution layer to aggregate information from adjacent nodes. Following this, the outputs are subjected to a ReLU activation and a normalization layer to update the node representations. Additionally, a residual connection is implemented at this stage of node convolution (as described in Equation 2 in Table1) to enhance the learning process before proceeding to the next graph learning iteration.

#### Edge Representation Refinement

Edge features are enhanced through advanced convolutional processes that integrate node and edge information. As depicted in Equation 3 in Table 1, the process begins with applying a 2D convolution to the edge embeddings, followed by summing this with the outputs from the graph convolution layer applied to the node embeddings. Specifically, the output from the node convolution block is processed through an outer sum block to produce a pairwise nucleotide representation. This representation is then combined with data from the adjacency matrix and edge embedding matrix to refine the edge representation for subsequent convolution operations on edges. The combined output is then fed into a sigmoid activation function to compute the edge gates. Finally, the edge embeddings are added back to the input embeddings through a residual connection, as outlined in Equation 4 in Table 1.

#### Integration of recurrent layers for RNA structure prediction

Subsequent to graph convolution processing, the edge embeddings are further fed through bidirectional ConvLSTM layers [61, 62] and one dilated 2D convolutional layer that predict the probability of pairwise nucleotide interactions with a sigmoid activation function.

#### Model Training

The training of the model initiates by predicting the probabilities of nucleotides forming base pairs given the RNA sequences in the training dataset. This is followed by calculating the loss between these predicted basepair probabilities and the reference base pairs derived from the RNA secondary structure. To address class imbalance in sequence-dependent secondary structure predictions, we employ a binary cross-entropy loss function with a dynamic weighting scheme. Specifically, the weight for the positive class is dynamically assigned based on the actual number of base pairs per RNA in each training round.

The losses from the loss functions are then backpropagated to update the model parameters. Model training is performed using the Adam optimizer with a default learning rate of 0.001 and a weight decay of 2.5e-4 [63]. Training typically concludes within 50 epochs, incorporating an early stopping mechanism that stops training if no improvement in loss is observed on the validation data.

Throughout the training process, strategies such as weight regularization, dropout, and batch normalization are extensively utilized to mitigate overfitting and ensure stable training dynamics. The deep learning model was trained on a machine equipped with an NVIDIA RTX A4000 GPU (16GB memory), while hyper-parameter tuning was extensively conducted on Nvidia A100 GPU (40GB memory) clusters provided by an NSF-funded cyberinfrastructure ecosystem, named ACCESS, which refers to the Advanced Cyberinfrastructure Coordination Ecosystem: Services & Support [64].

### 3.4. Model Evaluation

In this work, we use F1-score as a major metrics for evaluating the performance of different GNN algorithms for secondary structure generation. The F1-score is defined as

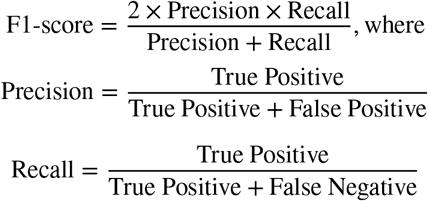

The True Positive in the evaluation of RNA secondary structure prediction refers to a correctly predicted base pair, i.e., a pair that is both in the predicted and the reference structure. A false positive, conversely, is a base pair that is predicted but does not exist in the reference structure. Similarly, a false negative represents a base pair that exists in the reference structure but is not predicted by the model. Thus, evaluating precision involves determining how many of the predicted base pairs are correct, while recall assesses how many of the actual base pairs were successfully predicted by the algorithm. The F1-score metric was calculated at the level of global RNA structure, with a higher score highlighting a more robust capability to deliver better secondary structure predictions.

## 4. Experiments and Analysis

In this section, we assess the performance of the proposed GNN methods compared to traditional energy-based methods, which provide initial edge connectivity for the GNN algorithms. We anticipated that the GNN algorithms would outperform, yielding more accurate predictions of RNA secondary structures.

### 4.1. Evaluation of GNN algorithms for RNA secondary structure prediction

We first compared the performance of six GNN methods against the established LinearPartition method for predicting RNA secondary structures, using F1-scores as the metric of assessment across validation and test datasets for bpRNA and PDB. As shown in Table 2, each GNN method outperformed the LinearPartition reference, which achieved baseline F1-scores of 63.59% and 64.10% on the bpRNA validation and test sets, respectively, and 65.59%, 70.57% and 74.27% on the PDB validation, test set, and test_hard set, respectively.

**Table 2.**
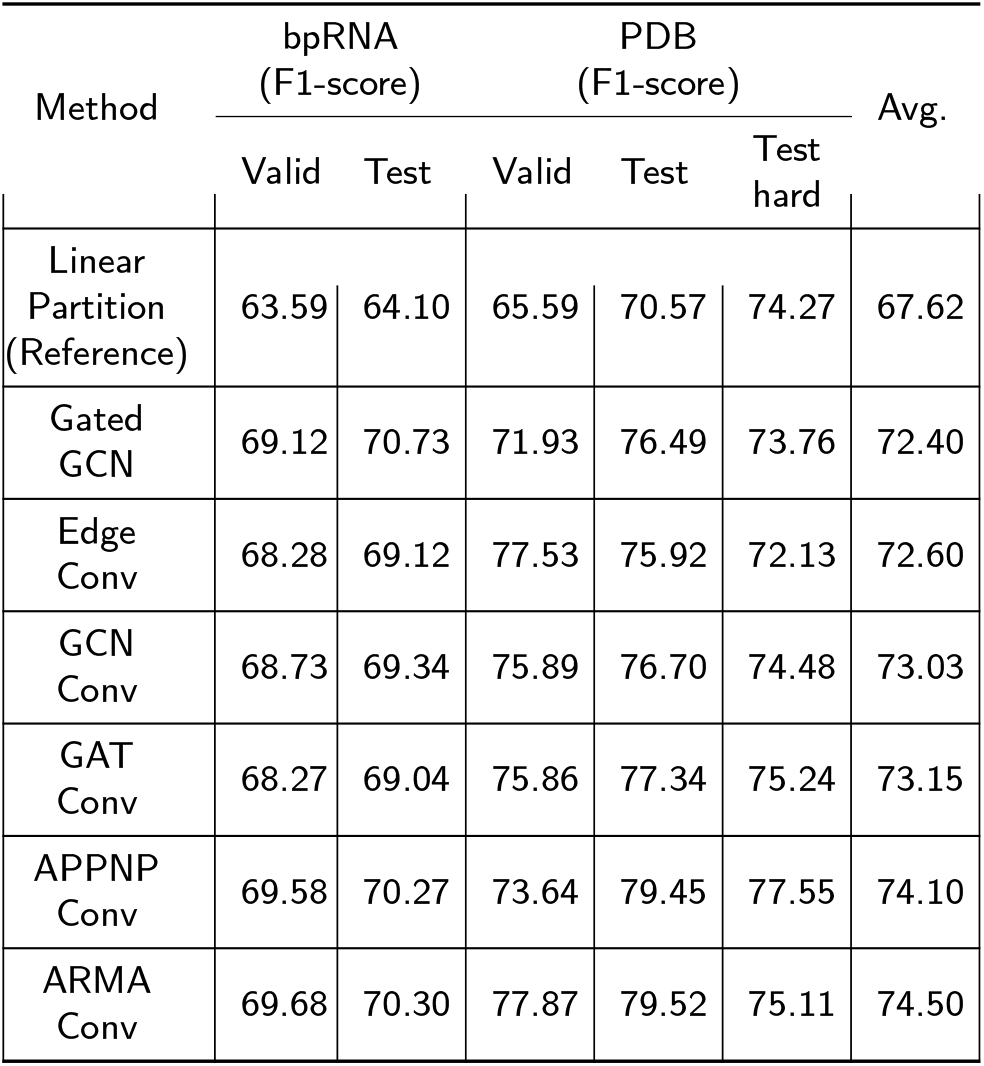
Performance evaluation of GNN methods in RNA secondary structure prediction using base-pair information from ‘Lin-earPartition’ as initial edge connectivity for GNN. The graph compares the F1-scores of various GNN methods against the energy-based method ‘LinearPartition’, which serves as the benchmark. All F1-scores are presented as percentages to quantify method’s performance.

| Method | bpRNA<br>(F1-score) |  | PDB<br>(F1-score) |  |  | Avg. |
| --- | --- | --- | --- | --- | --- | --- |
|  | Valid | Test | Valid | Test | Test hard |  |
| Linear Partition (Reference) | 63.59 | 64.10 | 65.59 | 70.57 | 74.27 | 67.62 |
| Gated GCN | 69.12 | 70.73 | 71.93 | 76.49 | 73.76 | 72.40 |
| Edge Conv | 68.28 | 69.12 | 77.53 | 75.92 | 72.13 | 72.60 |
| GCN Conv | 68.73 | 69.34 | 75.89 | 76.70 | 74.48 | 73.03 |
| GAT Conv | 68.27 | 69.04 | 75.86 | 77.34 | 75.24 | 73.15 |
| APPNP Conv | 69.58 | 70.27 | 73.64 | 79.45 | 77.55 | 74.10 |
| ARMA Conv | 69.68 | 70.30 | 77.87 | 79.52 | 75.11 | 74.50 |

Among the GNN approaches, GatedGCN achieved F1-score of 71.93% on validation and 76.49% on test sets, suggesting improved performance in capturing detailed relational patterns between nucleotides. The EdgeConv and GCNConv models also showed notable improvements, particularly on the PDB test set with F1-scores of 75.92% and 76.70%, respectively. Furthermore, the ARMAConv and APPNPConv methods, known for aggregating global contextual information, yield exceptional performance in the PDB test sets with scores of 79.52% and 79.52%, respectively.

Table 3 presents the performance of various GNN methods in predicting RNA secondary structures using ‘MXfold’ derived base-pair information for initializing graph edge connectivity. Across all evaluation sets, the F1-scores for these GNN models demonstrate consistent improvements. This analysis validated the effectiveness of utilizing refined edge connectivities, sourced from energy-based methods like LinearPartition and MXfold, to define the adjacency matrices for GNN inputs. The results support our hypothesis that integrating refined edge connectivity data into sophisticated GNN frameworks can lead to improvements over traditional secondary structure prediction methods.

**Table 3.** Performance evaluation of GNN methods in RNA secondary structure prediction using base-pair information from ‘MXfold’ as initial edge connectivity for GNN. The graph compares the F1-scores of various GNN methods against the energy-based method ‘MXfold’, which serves as the benchmark. All F1-scores are presented as percentages to quantify method’s performance.

| Method | bpRNA<br>(F1-score) |  | PDB<br>(F1-score) |  |  | Avg. |
| --- | --- | --- | --- | --- | --- | --- |
|  | Valid | Test | Valid | Test | Test hard |  |
| MXfold (Reference) | 64.59 | 65.68 | 77.90 | 77.15 | 79.30 | 72.92 |
| Gated GCN | 68.11 | 68.97 | 77.70 | 79.02 | 78.36 | 74.43 |
| Edge Conv | 65.65 | 66.77 | 76.67 | 77.92 | 76.56 | 72.71 |
| GCN Conv | 67.29 | 68.44 | 77.02 | 77.85 | 78.19 | 73.76 |
| GAT Conv | 66.70 | 67.39 | 75.98 | 78.46 | 79.68 | 73.64 |
| APPNP Conv | 66.91 | 67.81 | 77.01 | 78.00 | 77.38 | 73.42 |
| ARMA Conv | 67.03 | 68.09 | 78.24 | 79.20 | 78.33 | 74.18 |

### 4.2. Leveraging bpRNA and PDB Datasets for GNN model training

Furthermore, we investigated the effectiveness of graph neural network architectures for RNA secondary structure prediction in terms of three distinct training strategies: training exclusively on the bpRNA dataset, exclusively on the PDB dataset, and a hybrid approach involving pre-training on bpRNA followed by fine-tuning on PDB sets. The results reveal nuanced insights into how each training strategy impacts model performance across different datasets.

As shown in Figure 4 when models are exclusively trained on the bpRNA dataset, they demonstrate superior performance on the bpRNA validation and test sets with F1-scores significantly higher than those trained solely on PDB. For example, the GatedGCN model, when trained only on bpRNA, achieves an F1-score of 69.12% on the bpRNA validation, and 70.73% on the test, compared to 64.62% and 64.97%, respectively, when it is pre-trained on PDB only. However, the same bpRNA-trained models exhibit lower or comparable performance on the PDB dataset. Specifically, the APPNPConv method scores 67.27% on the PDB validation, 69.39% on the test, 64.74% on the test_hard set when trained only on bpRNA, notably lower than the 73.64%, 79.45%, 77.55% scores achieved after fine-tuning on PDB.

**Figure 4.**
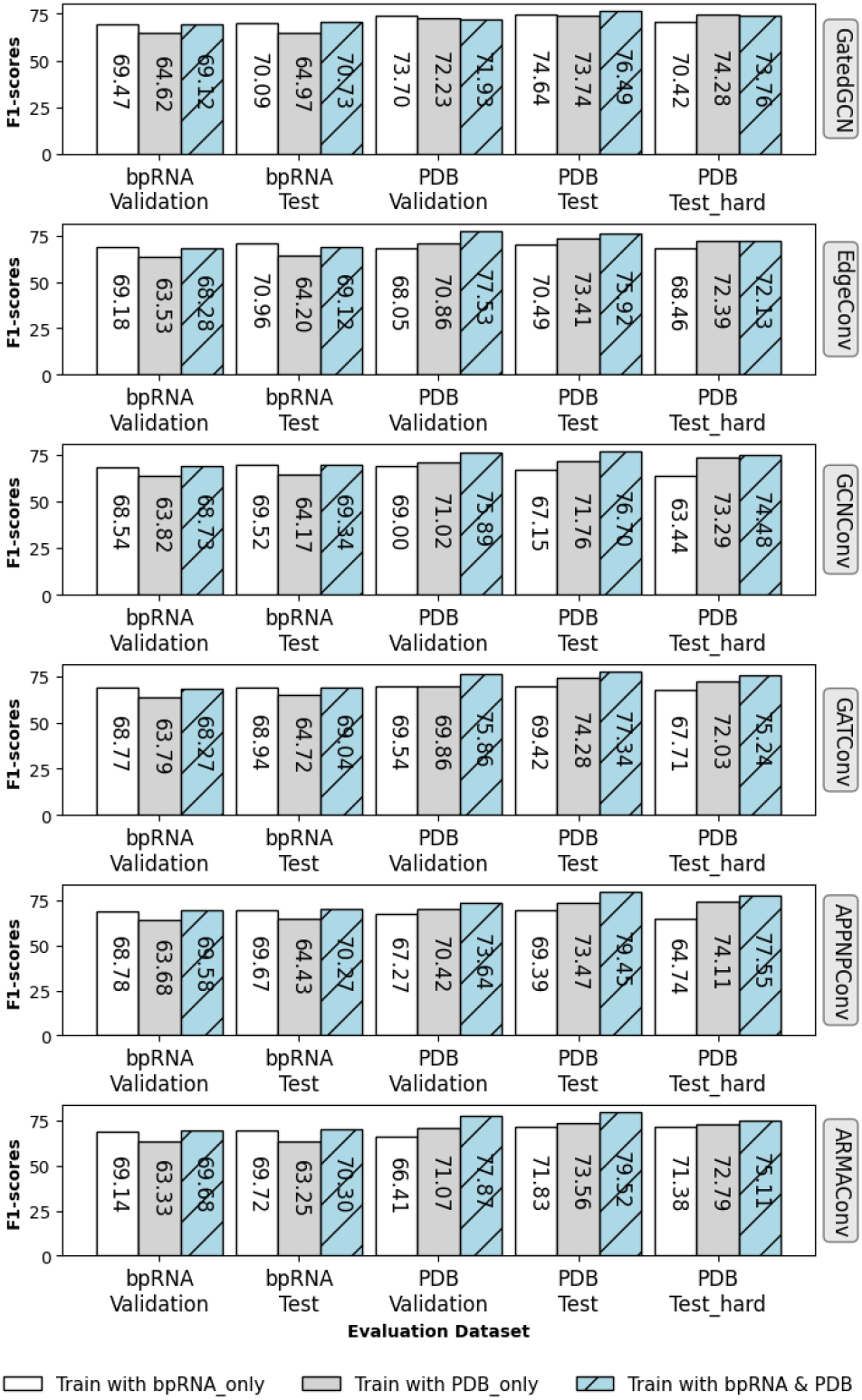
Comparision of F1-score on diverse datasets using the GNN models trained under three different experiments: (1) White Bar (Leftmost): Represents the F1-score when the GNN models were trained using only the bpRNA dataset; (2) Light Blue Bar (Middle): Represents the F1-score when the GNN models were trained using only the PDB dataset; (3) Striped Blue Bar (Rightmost): Represents the F1-score when the models were pre-trained on the bpRNA dataset and then fine-tuned on the PDB dataset.

Conversely, models trained exclusively on the PDB dataset show higher accuracy on PDB-related tests but underperform on the bpRNA dataset. For instance, the ARMAConv model scores significantly lower on the bpRNA test (63.25%) when trained only on PDB, compared to a higher score of 70.30% when the model is pre-trained on both bpRNA and PDB.

The hybrid training strategy, pre-training on bpRNA followed by fine-tuning on PDB dataset, results in the best performance across all datasets. This approach not only enhances the models’ accuracy on the PDB dataset but also maintains or improves their performance on the bpRNA dataset. Notably, the ARMAConv model achieves remarkable F1-scores of 77.89 on the PDB validation set, 79.52 on the test set, illustrating a significant improvement over its performance when trained on a single dataset.

These results highlight the benefits of pre-training GNN models with the bpRNA dataset to establish a robust baseline for RNA structure prediction. The performance is then effectively refined through subsequent fine-tuning on the PDB dataset, which leads to improved predictive accuracy and enhanced generalization capabilities.

### 4.3. Individual analysis for graph neural network over initial base pairs

The violin plot in Figure 5 provides a comprehensive comparison of the F1-scores based on predictions generated by two methods for 66 RNAs in PDB tests: LinearPartition and GNN (ARMAConv). From the visualization, we observe that GNN (ARMAConv) generally achieves higher F1-scores across the dataset, with a mean F1-score of 79.52% compared to 70.57% for LinearPartition. This suggests that GNN (ARMAConv) consistently outperforms LinearPartition in terms of average accuracy. It’s notable that while the upper quartile for LinearPartition (94.12%) is slightly higher compared to GNN (ARMAConv) (91.58%), the majority of its scores are concentrated below the mean, as indicated by a first quartile (Q1) of 49.09%, much lower than GNN (ARMAConv)’s Q1 of 75.37%. This lower Q1 suggests that a larger proportion of the F1-scores from LinearPartition are clustered at the lower scores.

**Figure 5.**
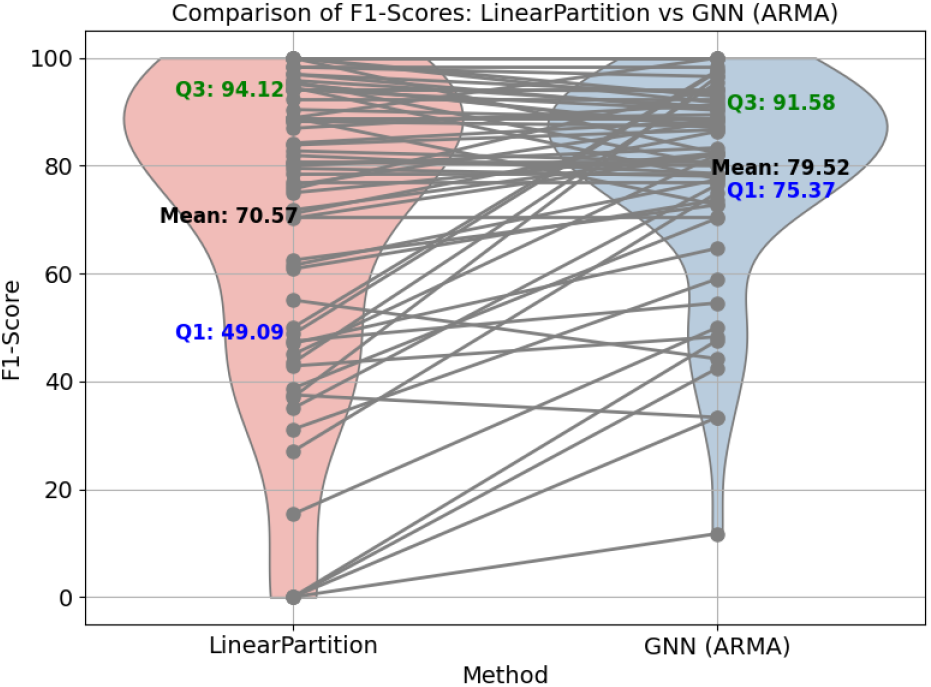
Comparison of F1-score on PDB test sets using methods LinearPartition and GNN (ARMAConv) for RNA secondary structure prediction.

The scatter plot shown in Figure 6 illustrates performance comparison between LinearPartition and GNN (ARMAConv) methods in predicting RNA secondary structures for 66 RNAs in PDB test set. The majority of the data points are located above the diagonal reference line, indicating that GNN (ARMAConv) typically achieves higher F1-scores than LinearPartition, with approximately 48.48% of the 66 targets showing improved F1-score under the GNN model. This suggests a superior efficacy of the GNN (ARMAConv) approach in handling a diverse range of RNA sequences. The statistical significance of this comparison is underscored by a p-value of 0.0003. While the bulk of the data clusters above the equality line, particularly at higher score values, there remain a few instances where LinearPartition matches or slightly exceeds the performance of GNN (ARMAConv). These cases might indicate specific sequence features or structural complexities where LinearPartition retains effectiveness, highlighting areas for potential refinement in the GNN approach. This comparative analysis demonstrates the robust capabilities of GNN (ARMAConv) in RNA secondary structure prediction, offering significant enhancements over traditional energy-based methods like LinearPartition. Figures 5 and 6 show cases where both the LinearPartition and GNN models underperform and instances where GNN surpasses LinearPartition models. Examining these structures can reveal edge-level features that could enhance prediction accuracy using GNN methods.

**Figure 6.**
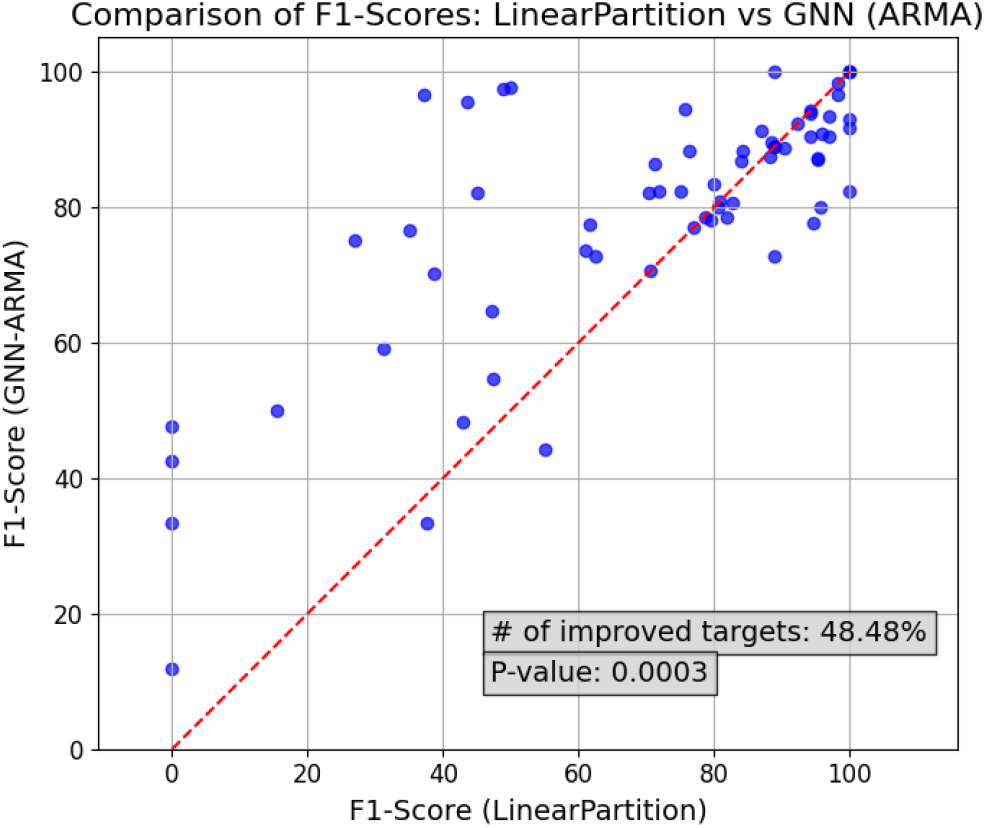
Comparative analysis of F1-scores in RNA secondary structure prediction on 66 RNA targets in PDB test dataset: GNN (ARMAConv) versus LinearPartition. Each point represents a single RNA from the PDB test set. The red dashed line is the diagonal reference line. Points above the line indicate instances where the GNN (ARMAConv) method achieved a higher F1-score than LinearPartition.

When utilizing base-pair predictions from ‘MXfold’ as the initial edge connectivity for the GNN (ARMAConv), the GNN model, as illustrated in Figure 7, achieved improved F1-scores for 39.39% of the 66 targets within the PDB test dataset.

**Figure 7.**
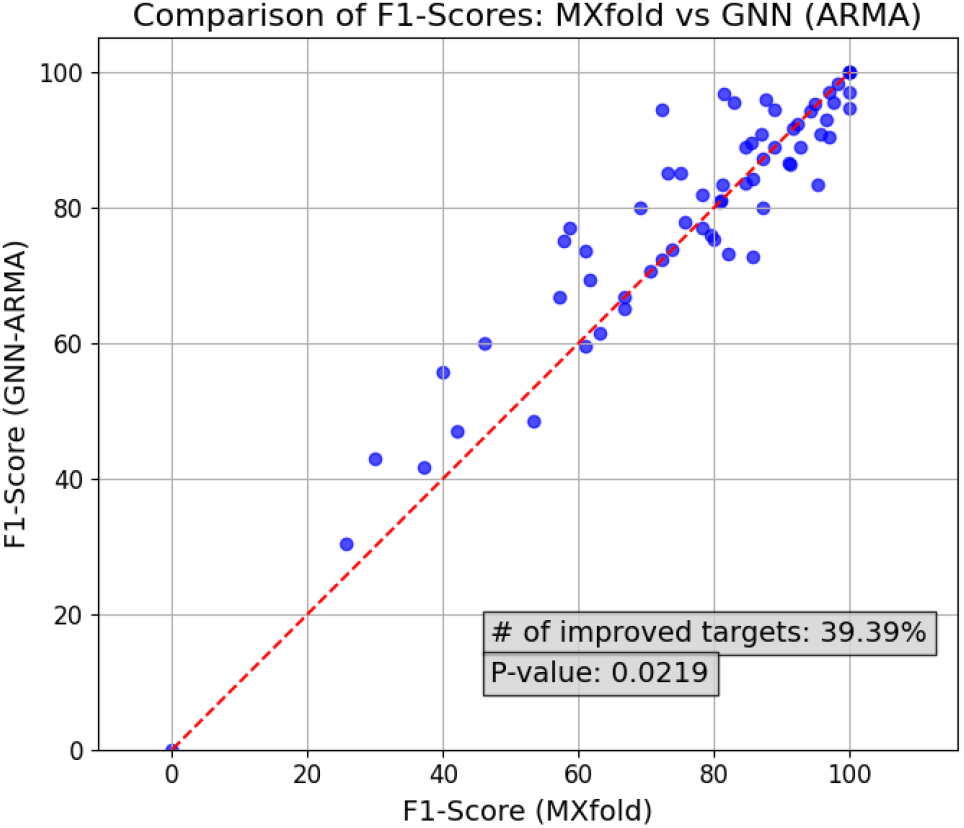
Comparative analysis of F1-scores in RNA secondary structure prediction on 66 RNA targets in PDB test dataset: GNN (ARMAConv) versus MXfold. Each point represents a single RNA from the PDB test set. The red dashed line is the diagonal reference line. Points above the line indicate instances where the GNN (ARMAConv) method achieved a higher F1-score than MXfold.

Figure 8 provides a comparative analysis of RNA secondary structure predictions made by LinearPartition, MX-Fold and two GNN models based on the ARMA architecture that use different edge adjacency matrices, against the native structure of an RNA sequence (PDB ID: 3AKZ, chain F). This structure was chosen because it contains a wide variety of RNA secondary structure motifs, including hairpin loops, internal loops, bulge loops, multiloop junctions, and double helical regions. The presence of these diverse motifs in a relatively compact structure facilitates visual comparison of different predictions. The native secondary structure was derived by characterizing the 3D structure using DSSR [65]. LinearPartition and MXfold achieve F1-scores of 43.48% (Figure 8 (B)) and 82.92% (Figure 8 (C)), indicating moderate accuracy with notable discrepancies in loop formations and base pairings. In contrast, the GNN (ARMAConv) models excel with a higher F1-score of 95.45%, closely mirroring the native structure and demonstrating its better capability to capture the intricate patterns and interactions within RNA sequences. This comparison highlights the potential improvement provided by GNNs in RNA secondary structure prediction over the free-energy based algorithms.

**Figure 8.**
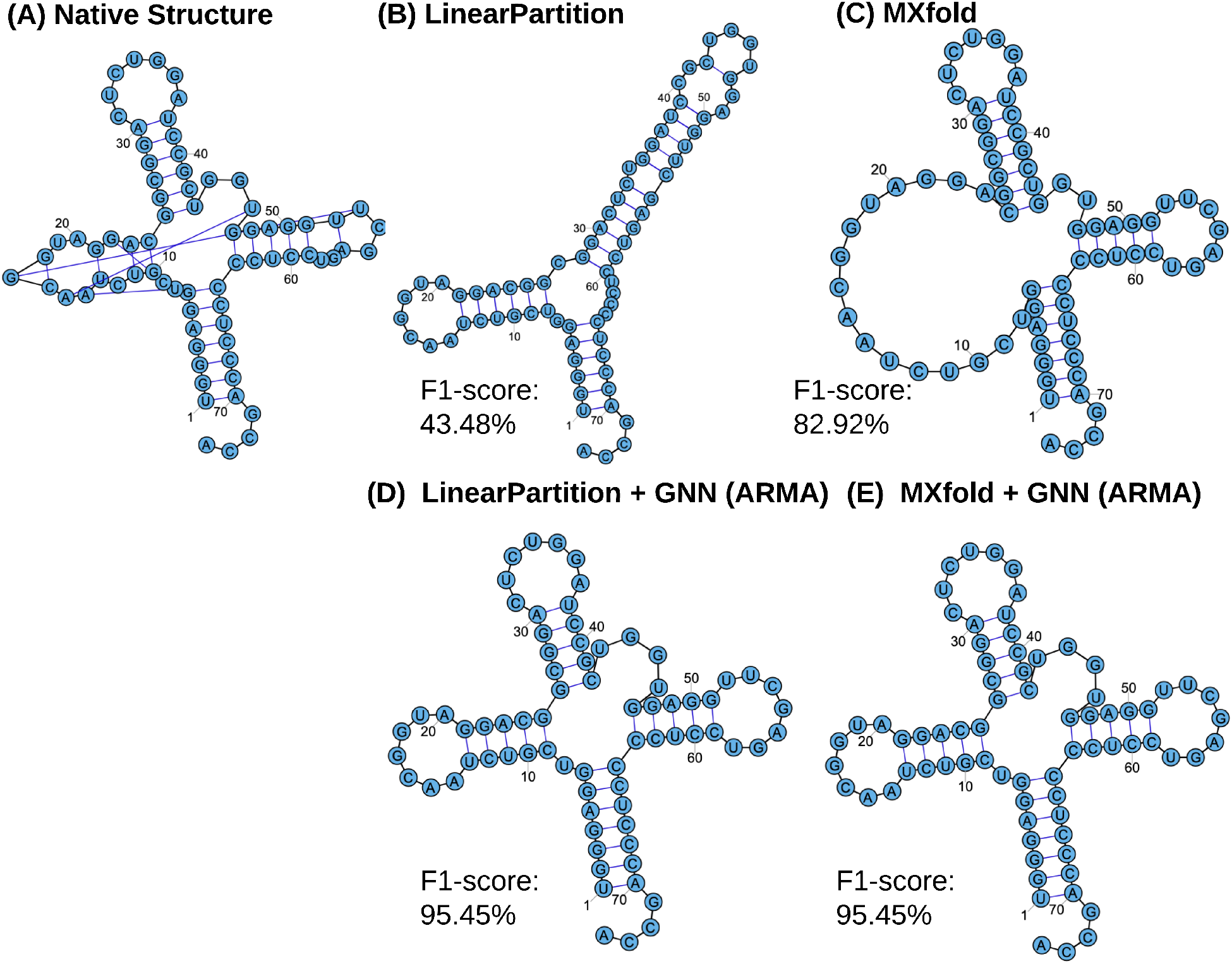
Comparative analysis of RNA secondary structure predictions for 3akz (Chain F). (A) displays the native structure, serving as the reference. (B) shows the structure as predicted by the method ‘LinearPartition.’ (C) presents the prediction from ‘MXfold.’ (D) depicts the structure predicted by GNN (ARMA) utilizing base-pair estimates derived from ‘LinearPartition.’ (E) reveals the prediction by GNN (ARMA) based on base-pair estimates from ‘MXfold.’

Specifically, from Figure 8, it is evident that all four predictive algorithms effectively avoid predicting the thermodynamically unstable single-nucleotide hairpin formed by a base pair between residues at positions 17 and 19 in the native structure, as illustrated in Figure 8 (A). However, none of the four predictive models managed to predict significant long-distance interactions between residues 18 and 50, 19 and 51, or 53 and 56, all of which are present in the native configuration. As shown in Figure 8 (B), LinearPartition tends to predict secondary structures by maximizing base pairing, potentially leading to highly stable but not necessarily biologically accurate representations of native folding. In contrast, MXfold’s predictions, as depicted in Figure 8 (B), often include extensive multi-junction loops that could contribute to thermodynamic instability. Our GNN models excel in accurately predicting these complex multi-junction loops at the structure’s core, as demonstrated in Figures 8 (C, D). This highlights the advantage of integrating GNN with outputs from other energy-based models to better capture the intricacies of native RNA folding without the bias towards overly stable configurations. By combining base pair probabilities from both LinearPartition and MXfold with GNN insights, we achieve a structure closely resembling the native one (as shown in Figures 8 (C, D)) and superior F1 scores relative to predictions based solely on energy-based methods, detailed in Table 4.

**Table 4.** Performance evaluation of GNN methods and traditional energy-based methods in RNA secondary structure prediction. All F1-scores are presented as percentages to quantify method’s performance.

| Method | bpRNA<br>(F1-score) |  | PDB<br>(F1-score) |  |  | Avg. |
| --- | --- | --- | --- | --- | --- | --- |
|  | Valid | Test | Valid | Test | Test hard |  |
| RNA fold [31] | 61.84 | 61.84 | 69.25 | 71.75 | 73.23 | 67.58 |
| Centroid fold [32] | 60.68 | 61.01 | 71.77 | 71.05 | 71.91 | 67.28 |
| Contra fold [33] | 64.72 | 65.33 | 72.63 | 74.36 | 74.87 | 70.38 |
| MXfold [35] | 64.59 | 65.68 | 77.90 | 77.15 | 79.30 | 72.92 |
| **MXfold + GNN (ARMA Conv)** | **67.03** | **68.09** | **78.24** | **79.20** | **78.33** | **74.18** |
| Linear Partition [34] | 63.59 | 64.10 | 65.59 | 70.57 | 74.27 | 67.62 |
| **Linear Partition + GNN (ARMA Conv)** | **69.68** | **70.30** | **77.87** | **79.52** | **75.11** | **74.50** |

### 4.4. Comparative performance evaluation of GNN methods and traditional energy-based methods

Table 4 provides a comparative performance analysis of the proposed GNN models alongside commonly used classical energy-based methods for RNA secondary structure prediction. Among the classical approaches, Contrafold displays commendable performance with an average F1-score of 70.38%. MXfold further advances this performance, yielding the highest scores among traditional methods with an average of 72.92% across all tests, achieving the best score at 79.30% in the PDB Test_hard dataset. However, the integration of GNNs with these traditional methods marks a significant improvement in predictive accuracy. MXfold combined with GNN (ARMAConv) leads to an average F1-score of 74.18%, demonstrating notable improvements across all tests, particularly with a 79.20% in the PDB Test. Similarly, LinearPartition, when integration with GNN (ARMAConv), exhibits a profound increase in performance, achieving an average score of 74.50%. These results underscore the efficacy of combining GNN architectures with traditional RNA folding algorithms to improve the accuracy of secondary structure prediction.

## 5. Conclusion

This study has demonstrated the potential of graph neural networks to advance the prediction of RNA secondary structures beyond the capabilities of traditional energy-based models. Through rigorous training and evaluations using both the bpRNA and PDB datasets, we have shown that GNNs not only outperform classical methods like LinearPartition but also provide a robust framework capable of capturing the complex interactions and long-range dependencies inherent in RNA sequences.

Our analyses across various GNN architectures underscore the effectiveness of integrating advanced graph-based learning in RNA structure modeling. The integration of pretraining on bpRNA datasets followed by fine-tuning on PDB datasets has proven particularly effective, yielding models that are well-generalized yet capable of achieving high accuracy in stringent test scenarios.

Furthermore, the improvement in prediction accuracies highlighted by our evaluations suggests that GNNs can be effectively tailored to enhance the RNA structural analysis beyond traditional energy-based methods.

Moving forward, we aim to refine our methodologies by incorporating additional motif features and diverse interaction types, such as predicting pseudoknots and other motifs beyond base-pairing, with the potential to further improve predictive accuracy. Additionally, we plan to expand this framework to encompass 3D RNA structure prediction. The effectiveness of language models in natural language processing for understanding RNA sequential patterns and evolutionary information has been well studied, demonstrating significant benefits in various RNA-related applications [66, 67, 68]. Given these successes, exploring the integration of RNA language models into GNN architectures to simultaneously predict both secondary and tertiary RNA structures presents a promising avenue for future research.

## 6. Acknowledgements

This work was supported by NIH Grant [2 R15 GM085699-04] awarded to B.M.Z. This work also used Indiana Jetstream2 GPU at Indiana University through allocation CIS240359 from the Advanced Cyberinfrastructure Coordination Ecosystem: Services & Support (ACCESS) program, which is supported by National Science Foundation grants 2138259, 2138286, 2138307, 2137603, and 2138296. We are grateful to Mr. Mahdi Rahbar, an MS-AI student at Saint Louis University, and Ms. Indira Vats, an undergraduate intern, for their invaluable support in exploring GNN architectures and conducting the literature review.

## 7. Code availability

The code and data associated with this study are publicly accessible at https://github.com/JieHou-SLU/RNASS. The package includes diverse scipts/notebooks to reproduce the GNN training, results evaluation, and demo codes to run traditional RNA secondary structure prediction tools.

## Notes

### Competing Interest Statement

The authors have declared no competing interest.

